# Unbiased co-occurring mutational landscapes in tumorigenesis

**DOI:** 10.1101/2025.01.19.633809

**Authors:** Yizhou Li, Yangfang Xiong, Wensheng Wei

## Abstract

Mutations with a synergistic effect on tumorigenesis were positively selected during tumor evolution, showing a tendency to co-occur in cancer genomes. To systematically gain insight of tumorigenic crosstalk at the mutation level, we utilized the FP-Growth algorithm in frequent pattern mining to create an unbiased map of co-occurring mutations from clinical genome sequencing data. We identified 100,933 frequent co-occurring mutation pairs across 22 cancer types. The co-mutation pairs involving cancer driver genes highlighted potential mechanisms for collaborative tumor formation, suggesting promising targets to disrupt tumorigenesis. Additionally, the large number of intra-gene co-mutations with strong dependencies indicated possible combinational effects on the oncogenic properties of specific proteins, which might be overlooked when individual mutations are not hotspots. Furthermore, our whole-genome mining strategy revealed a significant number of co-mutations in non-coding regions, with an enrichment of regulatory elements suggesting their potential role in modulating the co-expression of related genes. Finally, we demonstrated that our pipeline could systematically identify high-order co-mutations. In summary, our study provides the most extensive co-occurring mutations in tumor development, offering valuable insights into mechanisms and potential therapeutic targets.

## Introduction

The alteration in the cancer genome follows an evolutionary trajectory, where tumorigenic mutations continuously accumulate based on the current genomic context^1^. Early mutations selectively reshape the genome by retaining new mutations that confer a fitness advantage over the exiting landscape. Consequently, mutations that synergize in carcinogenesis are more likely to co-occur in the cancer genome, reflecting genetic epistasis in tumor evolution. Understanding this synergistic contribution is profoundly important for studying mechanisms and offers novel therapeutic opportunities.

Numerous studies have mined co-occurrence mutational patterns in genome sequencing data from clinical cancer patients, showing that these co-occurrence are widespread across diverse cancer types^2–4^. However, most previous analyses have focused on mutation events at the gene level, neglecting specific mutational types within gene bodies. Such strategy may raise several issues. Firstly, longer genes are more likely to harbor mutations, and studies have shown that genes suffer disparate tumor mutational burden (TMB)^1^ even when gene length is considered, increasing the false positives for co-occurrence in genes with significantly long lengths or low TMB. Additionally, large-scale functional studies revealed that mutations within the same gene can have various or even opposite effects, including large number of gain-of-function (GOF) mutations that behave independently of protein abundance^5,6^, as well as mutations that function dominantly-negative. These observations highlight that treating different mutations within the same gene equally is a crude approach to gain insight of synergetic signatures in the tumor genome.

It has been experimentally confirmed that tumorigenic synergy can arise from two individual somatic mutations, either mediated by missense or synonymous mutations in protein-coding regions^7,8^ or promoter mutations in non-coding regions^9^, both exhibiting significant co-occurrence patterns in clinical data. Despite some studies aiming to identify co-occurring mutations on a larger scale, the vast number of mutations detected in cancer patients makes it unaffordable to calculate co-occurrence frequency for each mutation pair. Therefore, previous endeavors have taken alternative approaches by selectively focusing on highly prevalent mutations or concentrating on mutations in specific pathways based on prior knowledge^10^. However, these results are limited by the original mutation dataset and mainly focused on coding regions. To the best of our knowledge, no study has yet extensively identified tumorigenic co-occurring mutations (co-mutations) in an unbiased manner.

To systematically predict mutation-mediated synergy in tumorigenesis, we developed a bioinformatics pipeline for massive identification of co-mutation pairs in cancer patients’ whole genome. Based on FP-Growth, an iterative frequent pattern mining algorithm^11^, we achieved an unbiased identification of highly prevalent mutation pairs and followed by certain filtering steps, we obtained 100,933 co-mutation pairs across 22 cancer types in the Internatinoal Cancer Genome Consortium (ICGC) database^12^. We interpreted the observed potential synergy effect from both inter-gene and intra-gene perspectives and highlighted the considerable proportion of co-mutations in non-coding regions. Finally, we demonstrated the pipeline’s capability of mining even high-order co-mutations. Our study provided an unprecedented predictive resource for understanding mutation-mediated synergy in tumorigenesis and potential novel therapeutic opportunities. Furthermore, we demonstrated the large-scale co-occurring mining pipeline is valuable for studying other biological questions. All information and results are presented on the website (https://co-mutation.streamlit.app/) to provide a user-friendly search tool.

## Results

### Unbiased co-occurring mutation mining in the ICGC database

We chose ICGC database as our original data source for the inclusion of massive coding mutaitons as well as non-coding mutations. The simple somatic mutation (SSM) dataset (STAR Methods) downloaded from ICGC contained 81,782,588 SSMs from 24,289 patients. Nearly half of the mutations occurred in intergenic regions (49.67%) and 45.16% in intronic regions. The coding mutations mainly included missense mutations, synonymous mutations, frameshift mutations and splice site-related mutations (Figure S1A).

The initial data preprocessing was mainly based on two considerations. First, we excluded mutations within annotated repeat regions according to RepeatMasker database. As multiple evidence demonstrated that the highly enriched mutations in repeated regions were created by error-prone repair or slippage during DNA replication^13^, we thus suspected the complicated mutational pattern in repeat regions might not reflect an evolutionary selection during tumor development. Moreover, we filtered the cancer types with less than 100 donors, for the lack of statistical power due to extreme small sample size. After the above procedures, our SSM dataset for downstream co-occurrence mining contained 36,624,683 SSM in 19,152 donors collectively, covering 45 cancer types (STAR Methods).

To address the heterogeneity of genetic alteration across different cancer types during tumor development, we constructed our mining pipeline by initially categorizing all donors according to their cancer types and performing the same workflow for each type in parallel (Figure 1A; STAR Methods). The FP-Growth algorithm^11^ took a mutation set for each donor as input and ended up with offering all mutation pairs with co-occurrence frequency higher than a defined minimum threshold. The mining procedures benefited from the construction of a FP-tree, a graph that effectively expressed the connectivity between mutations, to avoid pairwise frequency calculation through iteratively updated the FP-tree (STAR Methods). Considering two mutations co-occurred in a single donor contradicted the rationality because it could be any of the mutational combinations in any donors, we set the minimum threshold at 2, meaning each pair co-occurred in at least 2 donors for a given cancer type to achieve an unbiased mining for all of the reasonable co-occurring mutation pairs.

**Figure 1.**
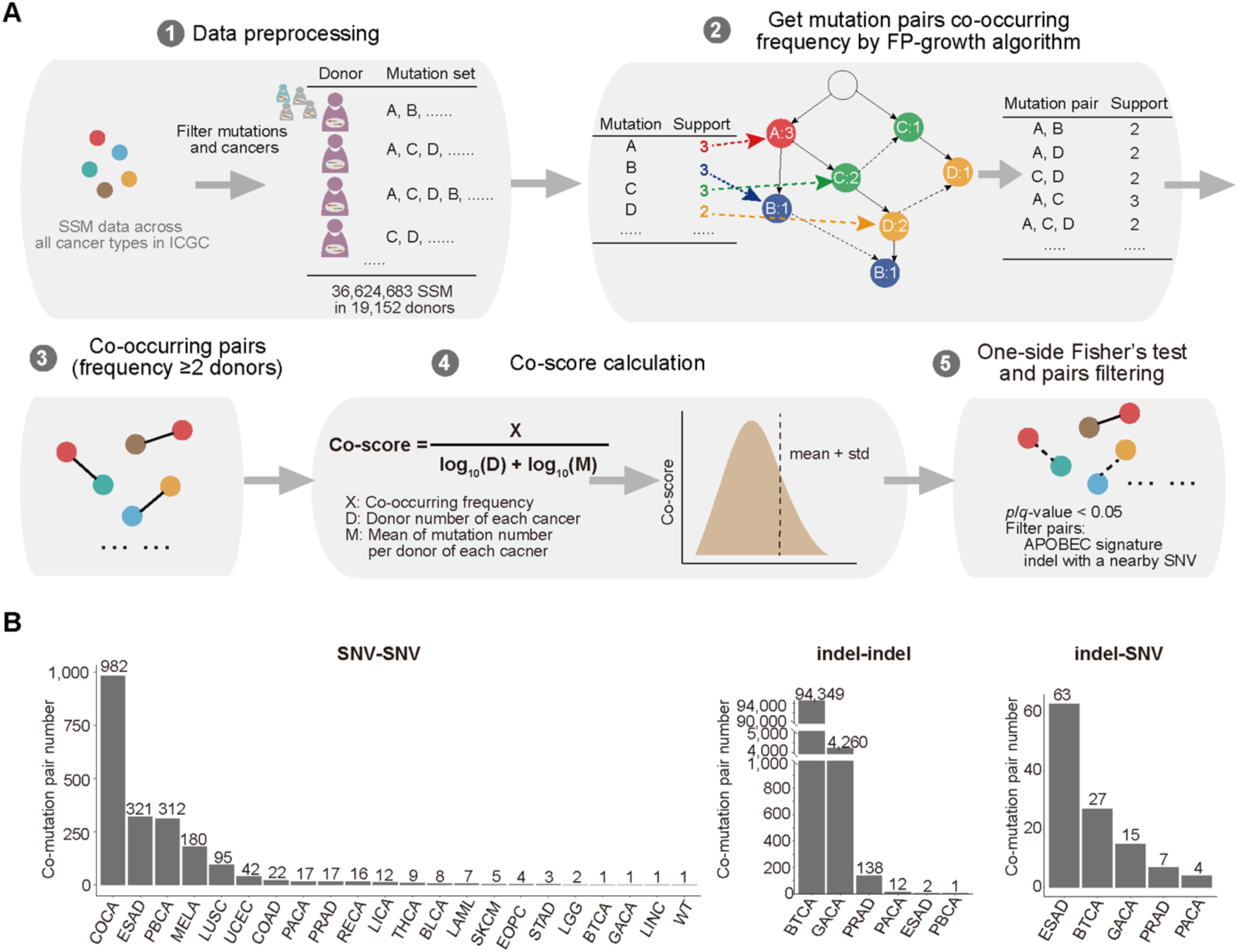
Systematically mining co-occurring mutation pairs in ICGC database. (A) Schematic of the computational workflow for unbiased large-scale mining of co-occurring mutation pairs in patients. (1) Preprocessing data from ICGC database. For each cancer type, mutations sets per donor is compiled. (2) Calculate the frequency of occurrence of mutation pairs by the FP-growth algorithm according to the conditions we set. (3) Get mutation pairs with co-occurring frequency larger than two donors. (4) Calculate co-score for each pair to evaluate co-occurring degree. (5) Filter pairs based on one-sided Fisher’s exact test and other two principles. (B) Mutation pair numbers of different cancer types. Mutation pairs are classified into SNV-SNV, indel-indel and indel-SNV based on the corresponding mutation types.

We next tented to determine a threshold to identify mutation pairs with more prevalent co-occrring frequency across all cancer types, as their unusual co-occurring pattern might be the outcomes of positive selection during tumor evolution. We reasoned the absolute count of donor number provided by FP-Growth roughly represented the co-occurring degree for each pair, but was simultaneously influenced by other two independent variations among cancer types. First, we noticed that the average mutation number carrying by a single donor varied significantly in different cancer types, reflecting the intrinsic diversity of mutational abundance in cancer type level (Figure S1B). In this scenario, the cancer types with richer mutational abundance were more likely to exhibit co-mutations under a given frequency threshold. Second, the cancer types with large sample size also would have an advantage in such analysis based on the absolute value of frequency (Figure S1C). To collectively assess the co-occurring degree of each pair in all cancer types, we introduced co-occurring score (co-score) as a normalized value to provide an unified evaluation of co-occurring degree for pairs. Basically, co-score took the variations of overall mutational abundance and donor number among cancer types into consideration, and served as a normalized value of raw donor count to make the co-occurring degree comparable between cancer types. According to the distribution of co-score (Figure S1D), we eventually set 0.97 (mean + std.) as threshold to identify prevalent co-mutation pairs (STAR Methods). To obtain a more reliable result, our workflow subsequently included several filtering steps. First, as it has been demonstrated that the co-occurrence frequency follows a hypergeometric distribution^14^, we conducted one-sided Fisher’s exact test followed by Benjamini-Hochberg adjustment to reduce the random combination of prevalent mutations (STAR Methods). Second, we filtered out pairs matched potential APOBEC mutational signature, as APOBEC might lead to some clustered mutation patterns such as kataegis and omikli^15^. Last, we also excluded pairs between an indel and a nearby single nucleotide variant (SNV), for these pairs were probably caused by variant miscalling (STAR Methods). Based on our workflow, the final results contained 100,933 co-mutations across 22 cancer types and we further classified them into three categories, which were SNV-SNV, indel-indel and indel-SNV due to the different evolutionary mechanisms of SNV and indel (Figure 1B; Table S1). In our observations, SNV-SNV covered 22 cancer types, which was much more prevalent compared with other two indel involved types, suggesting the synergy mediated by SNV might contribute to multiple cancer developments. On the other hand, indel-involved pairs were only found in 6 cancer types, but with significant number in biliary tract cancer (BTCA) and gastric cancer (GACA). We therefore inspected the compositions of mutations and co-mutation pairs in these two cancer types and observed that compared with SNV, indel still took the minortity part of the total SSM, which was consistent with other cancer types (Figure S1E). However, unlike SNV-SNV pairs which were mainly mediated by intergenic mutations, about one third of indel-indel pairs were between intronic mutations (Figure S1F), implying the instability of intron genome may mediate potential synergy in gene regulatory level, which might particularly contribute to the development of biliary tract cancer and gastric cancer.

### Co-mutation in cancer drivers suggests collaborative determination for tumor formation

Previous research has indicated that driver mutations may require assistance to stabilize or enhance the tumorigenic process through upstream or downstream interactions or by activating two collaborating oncogenic pathways^16^. We therefore suspected our results might contain such co-mutation pairs that were participated by single or double cancer driver genes. Indeed, we observed 8,625 (8.55%) paris from 20 cancer types involved mutations in established cancer driver genes according to Cancer Gene Census (CGC)^17^, with 4,081 pairs involved 139 oncogenes and 5,538 pairs involved 163 tumor suppressor genes (Figures 2A and 2B). 47 drivers paired across multiple cancer types, with the most extensive genes being *KRAS*, *KMT2C*, *MUC4*, *NF1* and *BRAF*, while 220 drivers showed cancer-specific pairings (Figure S2A). The considerable proportion of co-mutation pairs mediated by driver genes in various cancer types suggested the widespread collaborative determination for tumor formation by co-mutations.

**Figure 2.**
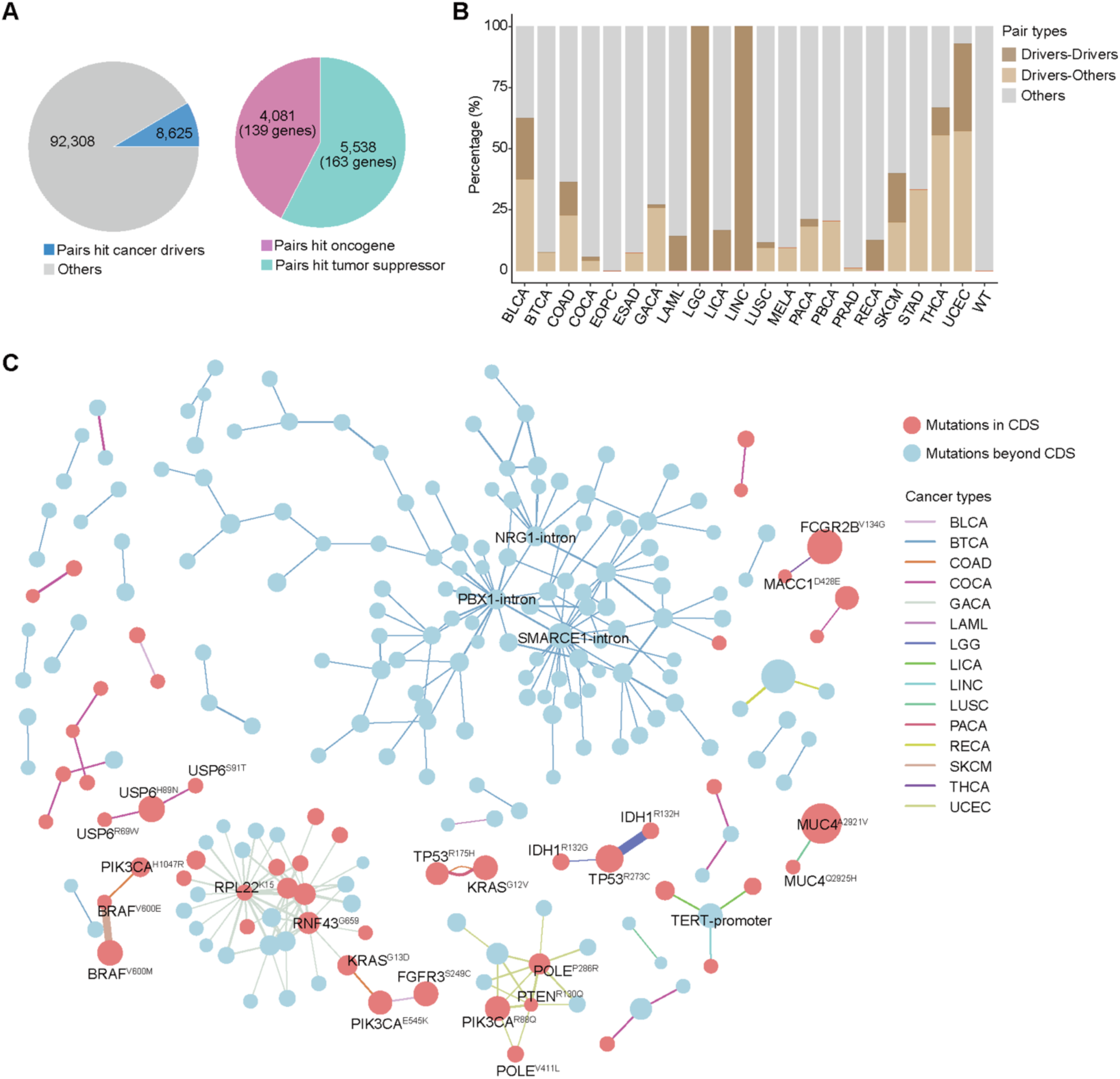
Inter-gene co-mutations involving cancer drivers indicated potential synergy for tumor early formation. (A) Pie chart showing the proportion of pairs involving known cancer drivers (left) and proportions of involving oncogenes and tumor suppressors (right). (B) Barplot showing the proportions of driver-involving pairs across cancer types. (C) Schematic showing the pairs between drivers. The pairs with co-occurring pattern are linked with the thickness of the line represented the absolute donor number carrying both mutations and the color represented various cancer types. The size of dot represents the individual prevalence of the mutation in corresponding cancer types. The dots colored in red represent coding mutations and blue represent non-coding mutations.

We next focused on the co-mutation pairs between known cancer drivers, for their tumorigenic roles were more explicit so far (Figure 2C). Generally, we observed the mutations in drivers paired with high cancer-type specificity. For example, a frameshift mutation in *RPL22*^K15^ showed high connectivity with other drivers, such as *RNF43*^G659^ and *UBR5*^K2120^, but exclusively in gastric cancer. Similar phenomenon was also occurred in *PIK3CA*^R88Q^, *POLE*^P286R^ and *PTEN*^R130Q^ in endometrial cancer (UCEC), indicating even though collaboration between drivers might be a shared mechanism in various cancers, but were mediated by specific players. However, very limited mutations participated in multiple cancers, such as *BRAF*^V600E^, *KRAS*^G12V^, *KRAS*^G13D^ and *PIK3CA*^E545K^.

We then investigated *KRAS* and *PIK3CA*, because they represented the most extensive participation in different cencer types. In pancreatic cancer, we observed a strong co-occurrence pattern between *TP53*^R175H^ and *KRAS*^G12V^. The synergistic mechanism between these two proteins has been well-documented, that *TP53*^R175H^ mutation can promote an isoform of the downstream factor GAP via the RNA binding protein hnRNPK, which in turn enhances RAS GTP levels to maintain KRAS’s abnormal activation and lead to stable oncogenesis in the pancreas^7^. Interestingly, while this mechanism was investigated using a *KRAS*^G12D^ mutant mouse model, only G12V showed a strong co-occurrence with *TP53*^R175H^ in the cohort we used, even though G12D was more prevalent individually. Notably, *TP53*^R175H^ with *KRAS*^G12V^ was shared in colorectal cancer (COAD) in our result and also aligned with another study on KRAS-associated mutations in NSCLC^18^, which found G12V had the highest co-occurrence frequency with *TP53* compared to other mutations. All of the observations suggested the tumorigenic synergy between *TP53*^R175H^ and *KRAS*^G12^ mutations might function in a pan-cancer manner, while G12V mediated the interaction more dominantly. Besides G12V, another prevalent participator was *KRAS*^G13D^, pairing with *RNF43*^G659^ in gastric cancer and *PIK3CA*^545K^ in colorectal cancer. Given the distinct effects of *KRAS* substitution^19^, our results suggested diverse underlying mechanisms of collaboration for different mutations mediated by *KRAS*.

For *PIK3CA*, we hit two hotspot mutations (E545K and H1047R) and a relatively rare mutation, R88Q. The two hotspot mutations all involved in colorectal cancer, pairing with *KRAS*^G13D^ and *BRAF*^V600E^ respectively, implying the overactivation of *PIK3CA* played a center role for colorectal cancer initiation. Interestingly, a rare mutation R88Q in adaptor-binding domain (ABD) showed high connectivity (pairing with *POLE*^P286R^, *POLE*^V411L^, *PTEN*^R130Q^ and *ARID1A*^R1989*^) and specificity in endometrial cancer. The mutations in ABD are considered as a strengthening for helical domain and kinase domain in *PIK3CA*^20^. However, in our result, none of the well-established hotspot mutations in helical or kinase domains were involved in endometrial cancer, neither were other detected mutations in ABD, suggesting a very unique crucial role of R88Q in endometrial cancer development.

The pair with strongest co-occurrence in our result was *IDH1*^R132H^ with *TP53*^R273C^ exclusively in glioma cancer (LGG), and notably, the other mutant, *IDH1*^R132C^ also paired with *TP53*^273C^ in the same cancer type, suggesting a highly specific selective preference between these mutations. The potential tumorigenic interaction between these two proteins was further supported by either high co-expression level in glioma patients (Figure S2B) and worse overall survival in glioma patients^21^, which collectively highlighted the urgency to clarify the synergetic mechanism between *IDH1*^R132^ and *TP53*^273C^. Moreover, we observed three hotspot intronic mutations within *PBX1*, *SMARCE1* and *NRG1*, respectively, mediated the majority of the inter-drivers mutation pairs, implying the essential role of these genes in biliary tract cancer.

Together, our method successfully identified known oncogenic synergy at the mutation level and revealed numerous unknown inter-gene pairs. We suspected the co-mutations involving cancer driver genes are particularly important as they represent new opportunities for early tumor intervention. As our understanding of cancer driver genes continues to grow, our results will offer more insights into their synergistic effects in the future.

### Intra-gene pairs indicate combinational effects on protein oncogenic properties

To facilitate the interpretation of the dependency between co-mutations in our result, we introduced Jaccard index (J), a universal method to assesss the co-occurrence pattern (Figure 3A; STAR Methods). According to the calculation, the pair with larger J indicated stronger dependency, regardless the individual prevalence. We surprisingly found that the pairs with high dependency tended to behave in an intra-gene manner significantly (Figure 3B). The combinational mutations are particularly relevant in protein engineering, where multiple mutations within a single protein sometimes jointly regulated a specific function, although the mechanisms behind this synergy are often unexplained^22^. We therefore hypothesized that the mutations co-occurring dependently within a protein might reflect a cooperative contribution for oncogenic function.

**Figure 3.**
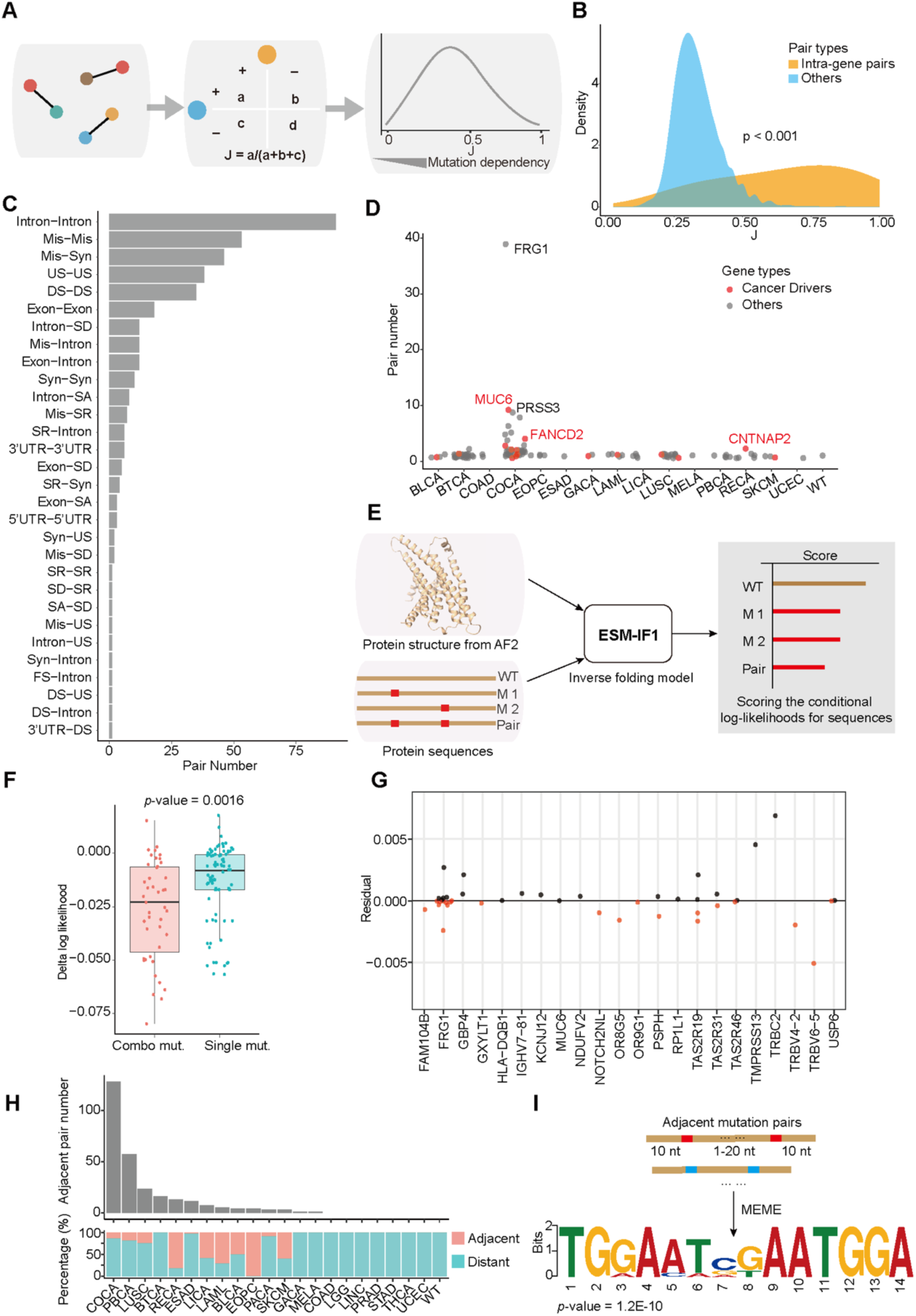
Intra-gene co-mutations with high dependency suggested combinational role in determing protein oncogenic function. (A) Workflow showing the Jaccard index calculation for each co-mutation. (B) Density plot showing the distribution of J in intra-gene pairs and others. The *p*-value was calculated by Wilcoxon test. (C) Histogram showing pair numbers of different intra-gene co-mtations classified by the specific mutation consequences within a same gene. Mis: missense mutation, Syn: synonymous mutation, US: gene upstream variant, DS: gene downstream variant, SD: splice donor site variant, SA: splice acceptor site variant, SR: splice region variant, FS: frameshift variant. (D) Scatter plot showing the distribution of intra-gene pairs number across cancer types. Each dot represents a given gene identified in the corresponding cancer types. The dots colored in red represent cancer drivers in CGC. (E) Workflow for evaluating the impact of missense mutations on protein structural stability using the ESM-IF1 model. The input files include the wild-type protein structure and the protein sequences for the wild-type, single mutants, and combinational mutant. The model calculates the log-likelihood for each sequence, with smaller values indicating greater destabilizing effect on protein stability. (F) Boxplot showing the effects of single and combinational mutations on protein stability. The delta log-likelihood represents the difference between the scores for the mutant sequence and the wild-type. Student’s t test, *p*-value = 0.0016. (G) Synergistic effect of mutation pairs on protein stability. The synergistic effect score for the double mutation was calculated by subtracting the sum of the delta log-likelihood for the two single mutations from the delta log-likelihood of the combinational mutation. The dots colored in red represent the synergistic effect scores less than 0, indicating the mutation pairs have synergistic effects in destabilizing the protein. (H) The number and percentage of adjacent mutation pairs across cancer types. Mutation pairs located within 20 nucleotides on the chromosome were defined as adjacent mutation pairs, and those with greater distance were distant mutation pairs. (I) Surrounding sequence motif of adjacent mutation pairs. We extended chromosomal positions by 10 base pairs on both sides of adjacent mutation pairs, extracted the corresponding sequences, and used MEME to de novo search for potential motifs.

To explore this, we first classified the intra-gene pairs by their consequence types (Figure 3C). A significant proportion of these mutations were in non-coding regions of the gene body (intron, gene downstream, gene upstream, and UTR), likely affecting cis-regulatory elements. The most prevalent coding intra-gene pairs were two missense mutations and one missense mutation with one nearby synonymous mutation. Notably, a similar phenomenon has been noticed and clarified in a *KRAS* co-mutation pair (G60G and Q61K) that caused an abnormal splicing event, leading to a novel drug-resistance phenotype^8^.

We then investigated the genes involved in intra-gene pairs. Overall, we observed 115 genes with intra-gene pairs in 16 cancer types, with 15 genes were known drivers genes, representing potential combinational effects for oncogenic function determinations (Figure 3D). Colorectal cancer proteins typically harbored more intensive intra-gene co-mutations compared to other cancer types, with *FRG1* being the most significant one containing 39 pairs, and 24 of them were between coding mutations rather than the more prevalent intronic pairs (Figure S3A). As an RNA binding protein, *FRG1* is part of the human spliceosome complex, stabilizing the active conformation of the human C complex^23^, the abnormally low expression of *FRG1* in colorectal cancer has been noticed^24^ while the mechanism remains unclear. We therefore hypothesized the intensive co-mutations might impact the protein fitness of *FRG1*, which further contribute to colorectal cancer development with tumor suppressor property.

To reinforce this idea, we collected all of the missense mutation mediated intra-genes pairs in our results and assisted from protein inverse folding prediction to achieve a large-scale evaluation of combinational impact on protein fitness, as a novel perspective to identify potential more tumor suppressors. By providing a wild-type structure information along with the mutant sequences to ESM-IF1^25^, the AI based model would evaluate a delta log-likelihood which indicated the likelihood that a given sequence can fold into the functional structure after undergoing specific mutations (Figure 3E; STAR Methods). Overall, we observed combinational mutations had more detrimental effect for protein fitness than single mutation for all 22 proteins (Figure 3F) and 21 pairs in 13 proteins with severer impact compared with simply adding the effect by two individual mutations, including *FRG1* mentioned above (Figure 3G). Interestingly, we found proteins of two families, olfactory receptors (ORs) and taste 2 receptors (TAS2Rs) were highly involved in the analysis, and also with most significant preference for intra-gene co-mutations when taking gene length into consideration (Figure S3B). The high selectivity indicated their tumor suppressor role caused by combinational mutations might be shared in certain protein families.

Moreover, we expanded the idea of combinational effect to DNA level and questioned if any adjacent co-mutations jointly destruct or create functional motifs in genome, which might further impact DNA binding event. We therefore classified all co-mutations within 20 bp by mutation position and performed de novo motif discovery analysis on the +/-10 bp sequences to identify sequence preferences of adjacent co-mutations (Figure 3H; STAR Methods). The analysis revealed a significant motif pattern (Figure 3I), showing high binding preference for ZNF671, a protein with well-knowledged tumor suppressor activity^26,27^, suggesting that this type of co-mutation pairs might function by disrupting ZNF671 binding motifs, contributing to tumorigenesis.

### The enrichment of non-coding mutations in regulatory elements suggests potential synergy through gene co-expression

Non-coding mutations accounted for the majority of the co-mutation pairs we identified (Figure 4A). Understanding the significance of non-coding mutations in tumor genome has been a long-standing challenge in this field. Therefore, we investigated whether our results could provide insights into the function of widespread non-coding mutations, especially those within regulatory elements.

**Figure 4.**
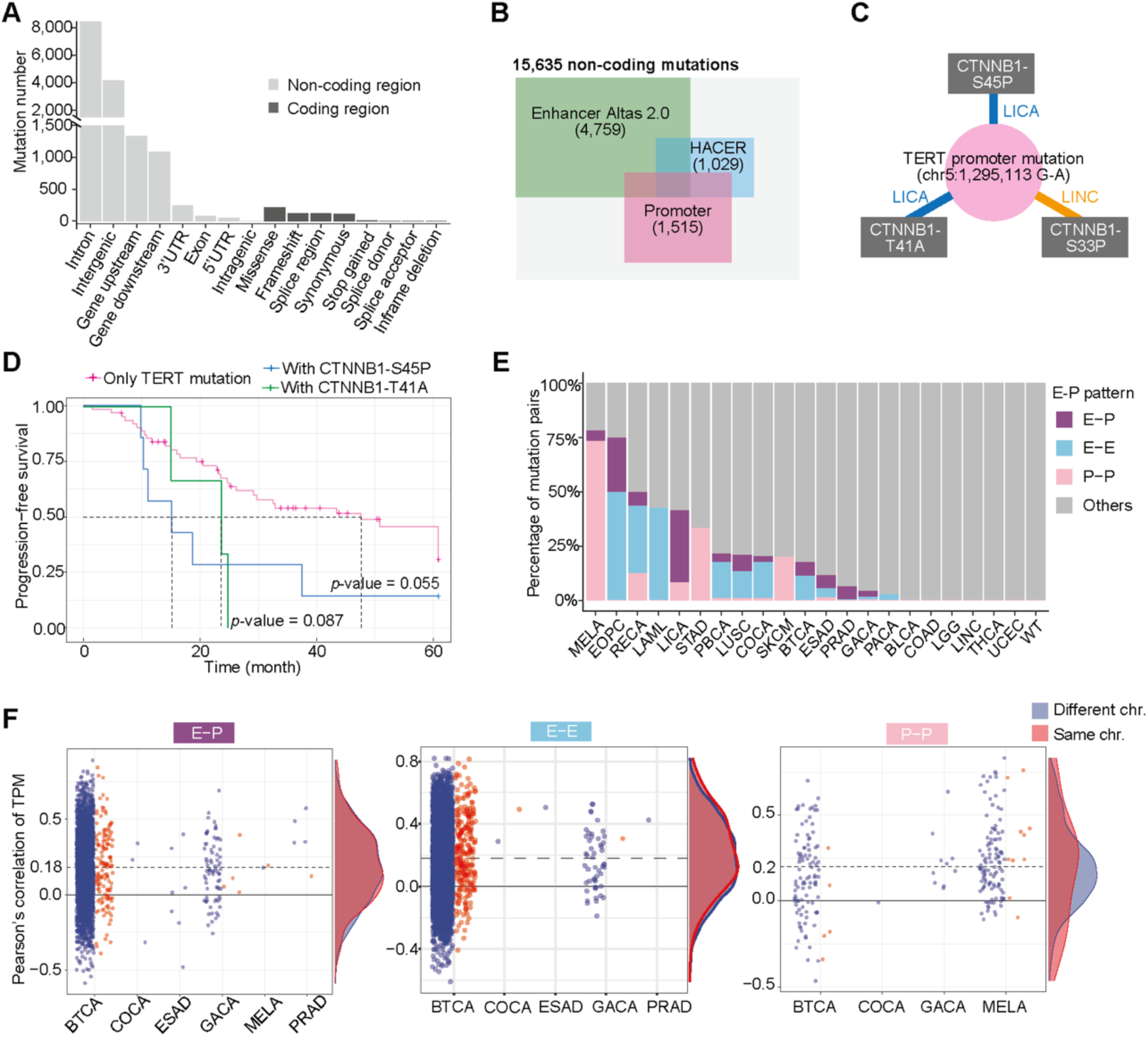
Non-coding pairs enriched in regulatory elements might represent unknown expression dependent functional crosstalk between two genes. (A) Histogram showing the consequence types of all mutations participated in our co-mutation results. (B) The number of non-coding mutations in enhancer region and promotor region in our results. (C) Schematic showing a specific *TERT* promoter mutation co-occurrs with different mutations of *CTNNB1* in LICA and LINC. (D) The comparision of progression-free survival rate of patients with only identified *TERT* promoter mutation and whom with both *TERT* and paired *CTNNB1* mutation in LICA. (E) Proportions of regulatory element involving mutation pairs across various cancers. The mutation pairs are classified by E-P (enhancer mutation with promoter mutation), P-P (promoter mutation with promoter mutation) and E-E (enhancer mutation with enhancer mutation). (F) Scatter plot showing the correlations between mRNA expression of gene pairs with E-P (left), E-E (middle), P-P (right) in tumor donors. Each dot in the plot represents a gene pair. The y-axis represents the Pearson’s correlation of the two genes in corresponding cancer types. The dashed line indicates the mean correlation of all gene pairs. The density distributions of the correlation coefficients are shown on the right. The different colors represent whether the mutation pairs are located on the same chromosome.

To explore this, we cross-referenced the non-coding mutations from our results with enhancer regions from the EnhancerAtlas 2.0 database and HACER database, and also promoter regions defined as 2,000 bp upstream of the annotated transcription start site (TSS) in the whole genome (Figure S4A; STAR Methods). We found that about one-third of non-coding mutations were located in enhancer regions and 1,515 mutations were found in promoter regions, suggesting our results were highly relevant to regulatory function (Figure 4B). For example, we observed a specific *TERT* promoter mutation co-occurring with three separate missense mutations in *CTNNB1*, a pattern observed exclusively in liver cancer but varying across different subtypes (Figure 4C). The combinations of this *TERT* promoter mutation and *CTNNB1* mutations were also clinically associated with worse survival rates in patients carrying both *CTNNB1*^S45P^ and *CTNNB1*^T41A^ (Figure 4D). Although the protein physical association between TERT and CTNNB1 has been previously detected^28^, the functional interaction remains unclear. Our results may be valuable for further mechanistic studies between these two genes. Additionally, numerous enhancer or promoter mutations with multiple combinations were identified, some of which were particularly significant, such as a promoter mutation of *RPL13A* (Figure S4B), implying its crucial role in melanoma development.

The functional interaction between regulatory elements could be mediated by either the physical contact formed by the 3D structure of chromosomes^29^ or other mechanisms acting in trans, such as enhancer RNA affecting downstream promoters, both of which can regulate gene expression^30^. Given that many of the non-coding mutations we identified were in enhancer and promotor regions, we focused on co-mutation pairs involving enhancer-enhancer (E-E), enhancer-promoter (E-P), and promoter-promoter (P-P) regions, hypothesizing that some could modulate unknown expression dependencies between two genes (Figures 4E and S4C).

We therefore systematically investigated gene co-expression level for the three types of pairs in corresponding cancer patients (STAR Methods). Overall, we observed 413 gene pairs related to E-E mutation pairs, 269 gene pairs related to E-P mutation pairs and 28 gene pairs related to P-P mutation pairs showed high co-expresssion level (Pearson’s correlation >= 0.5), indicating a large-scale identification of tumorigenic co-expression mediated by non-coding mutations in regulatory elements (Figure 4F).

### Mining high-order co-occurring mutations

So far, we have systematically analyzed the dependency between two mutations in cancer patients, but higher-order epistatic effects may also exist. One common method to study complex genetic interactions involves using yeast as experimental material and measuring the fitness of mutants to evaluate the epistatic effect between mutations. In a previous study on high-order interactions in yeast tRNA mutations, it was found that not only were second-order interactions abundant, but third-order and higher-order interactions were also widely present^31^. In addition, another study analyzing triple mutation effects in yeast found that three-gene interactions often occurred between functionally related genes, with third-order interactions more likely to bridge biological processes in the cell, despite having overall weaker effects than second-order interactions^32^.

To globally identify high-order mutational interactions in the human genome through fitness experiments is extremely challenging. However, we demonstrated that our method is particularly suited for mining high-order co-occurring mutations. By simply resetting the frequent pattern length in FP-Growth, we fine-tuned our pipeline to predict third-order mutational interactions in tumorigenesis systematically (Figures 5A and S5A). We eventually identified 18,373 third-order pairs in 13 cancer types (Figure 5B; Table S2) and observed the mutations participated in third-order pairs with strong overlap with second-order pairs (Figure 5C). For example, there were notable connections in gastric cancer centered on a frameshift mutation of *RPL22* and in colorectal cancer, some mutations identified in second-order pairs of *FRG1* were connected in an even higher-order manner (Figure S5B). All together, we highlighted that our pipeline can easily expand to predict high-order interactions, providing valuable results for further exploration.

**Figure 5.**
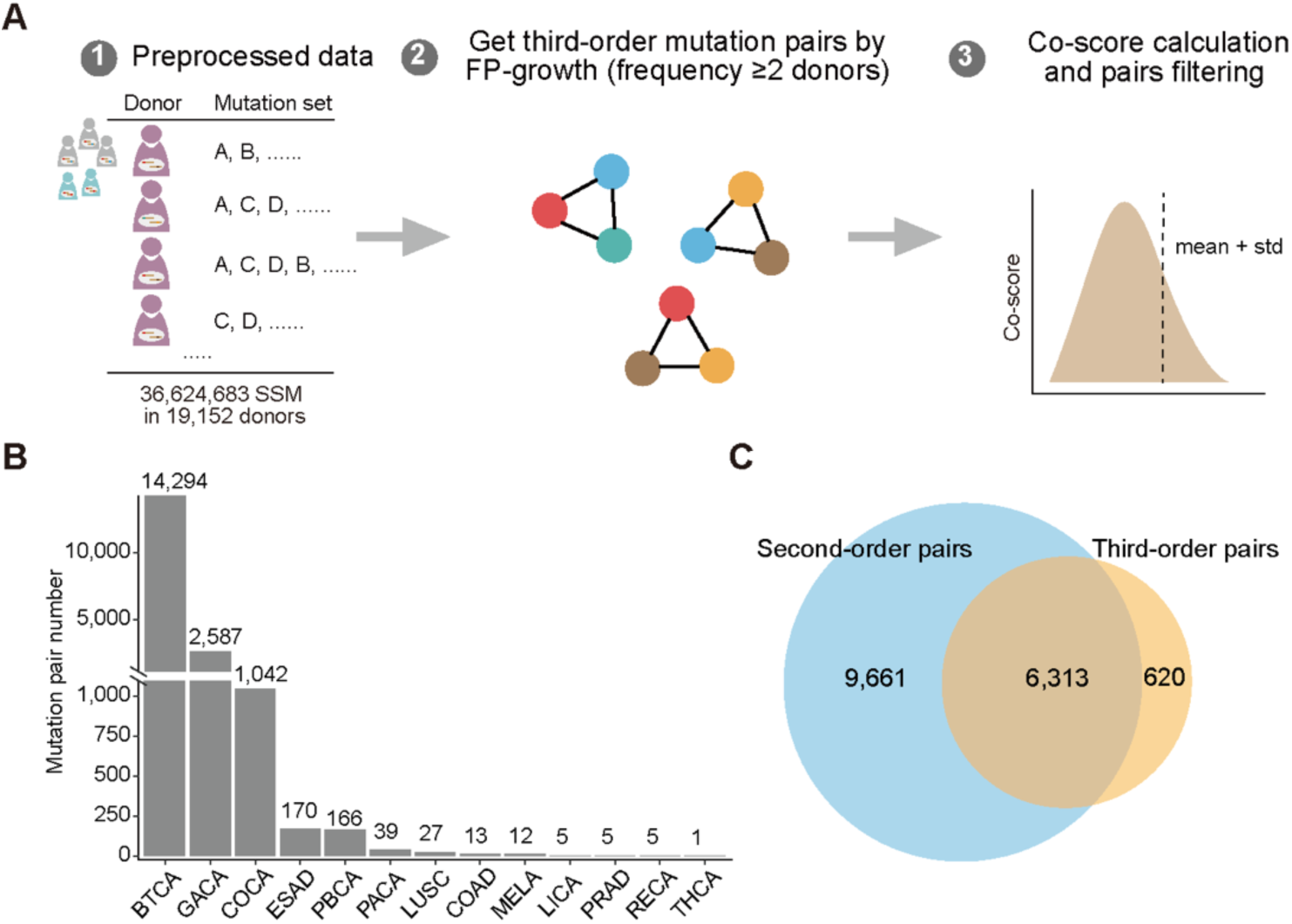
Schematic of workflow for high-order co-occurring mutation pairs mining. (A) Schematic of the computational workflow for third-order co-occurring mutation pairs mining. The filtered SSM dataset is as same as the mining of second-order mutation pairs. We set the length of the mutation pair as 3, then calculated the co-score and selected mutation pairs with a co-score above threshold, and filtered potential false positive mutation pairs. (B) The number of third-order mutation pairs across different cancers. (C) The number of overlapping mutations involved in second-order mutation pairs and third-order mutation pairs.

## Discussion

In this study, we took advantage of FP-Growth algorithm from the data mining filed to uncover an unbiased landscape of co-occurring mutations from ICGC database, suggesting mutation-mediated synergy in tumorigenesis. Given the complexity of tumor development and the significant lack of understanding, our investigation provides potential interaction clues at the mutation level and offers a novel perspective to interpret numerous mutations with unknown significance. Although many co-occurrence patterns cannot currently be explained mechanistically, we believe further experimental validation, especially large-scale functional screening assays, are necessary for more reliable identification.

We acknowledge that a major limitation of our current pipeline is the use of bulk sequencing data, meaning the observed co-mutations are carried by individual patients but may not be present in the same cell. In this scenario, the two mutations might function independently in tumor development without cellular crosstalk, potentially leading to potential false positives in our analysis. A more precise approach would involve clonal information or even single-cell sequencing data to perform a similar mining pipeline while considering cell heterogeneity. However, due to the lack of clonal information in the dataset we used and the large-scale, unified single-cell data across all cancer types, we chose bulk data to achieve a relatively comprehensive co-occurring mutational pattern, which already included many reasonable candidates.

In conclusion, the unbiased co-occurring mutational landscape identified in this study provides novel clues for understanding tumor development mechanisms. Additionally, disrupting these interactions could have therapeutic relevance, as interfering with either mutation could potentially block tumorigenesis and this rationale is particularly useful when one of the mutant proteins is difficult to target with drugs.

**Figure S1.**
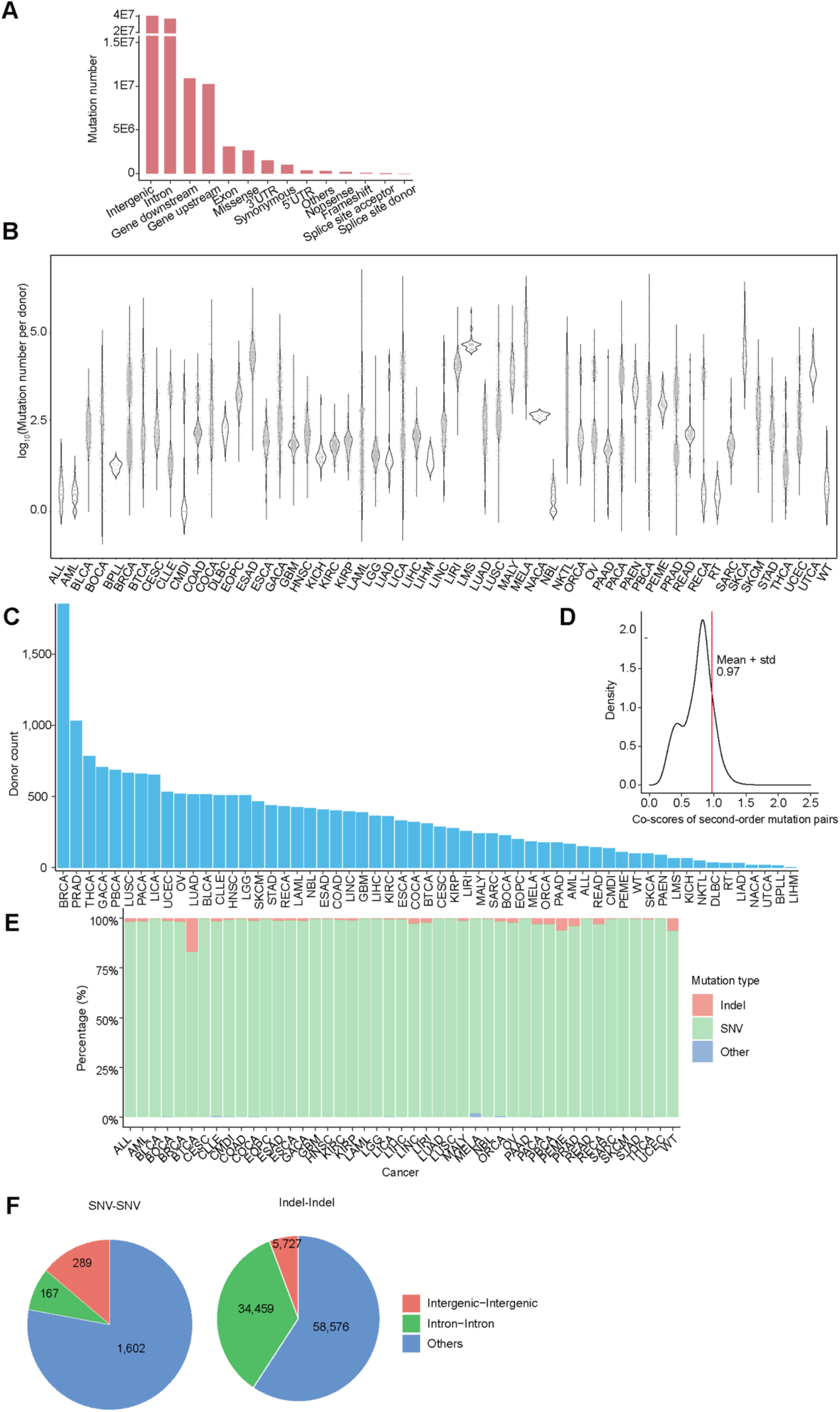
Mutation information in ICGC. (A) Barplot showing distribution of mutation types in ICGC database. (B) Variation in mutation numbers per donor across different cancer types in ICGC database. Each point means the log_10_(mutation numbers) of an individual donor. (C) The number of cancers donors in ICGC databases. Blue represents cancers with co-occurring mutation pairs in our results, while red represents cancers without co-occurring mutation pairs. (D) The co-score density of second-order mutation pairs. The mean plus standard deviation was about 0.97. To enhance the clarity of the image, the score range was set from 0 to 2.5, as there were only 144 mutation pairs with scores higher than 2.5. (E) The proportions of different mutation types in various cancers from the input SSM data. “SNV” refers to single nucleotide variations, “Indel” represents insertions or deletions, and “Others” refers to irregular base segment substitutions with a length greater than 1. (F) Pie charts showing the proportions of pairs between intergenic mutations and intronic mutations in SNV-SNV (left) and indel-indel (right).

**Figure S2.**
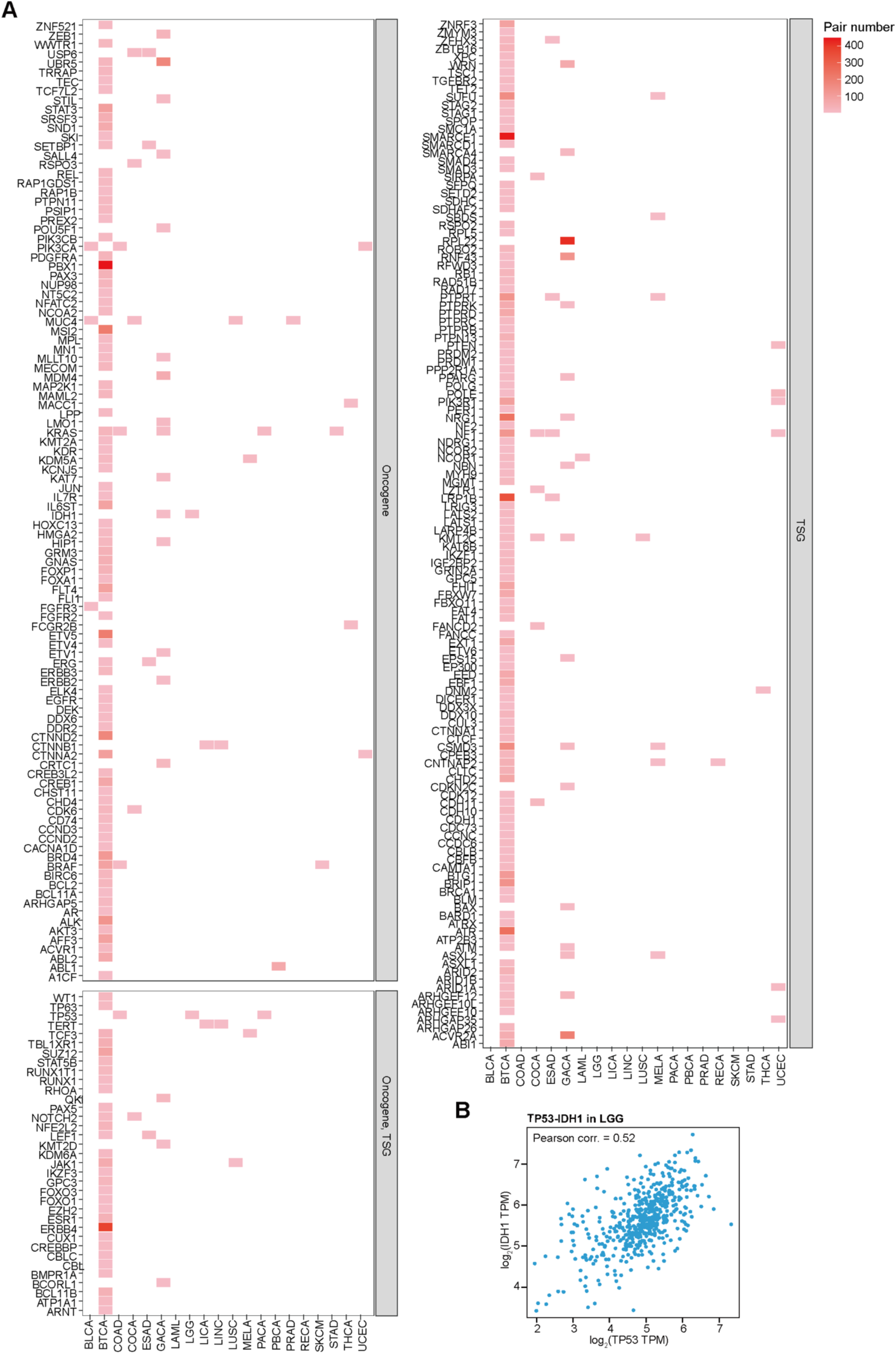
Pairs involving cancer drivers. (A) Heatmap showing cancer drivers with mutation pairs across cancer types, classified by oncogenes and tumor suppressor genes. Classifications of cancer drivers are from Cancer Gene Census database. TSG: Tumor suppressor gene. (B) Scatter plots showing the correlation of gene expression between *TP53* and *IDH1* in glioma patients. Each dot represents a glioma donor.

**Figure S3.**
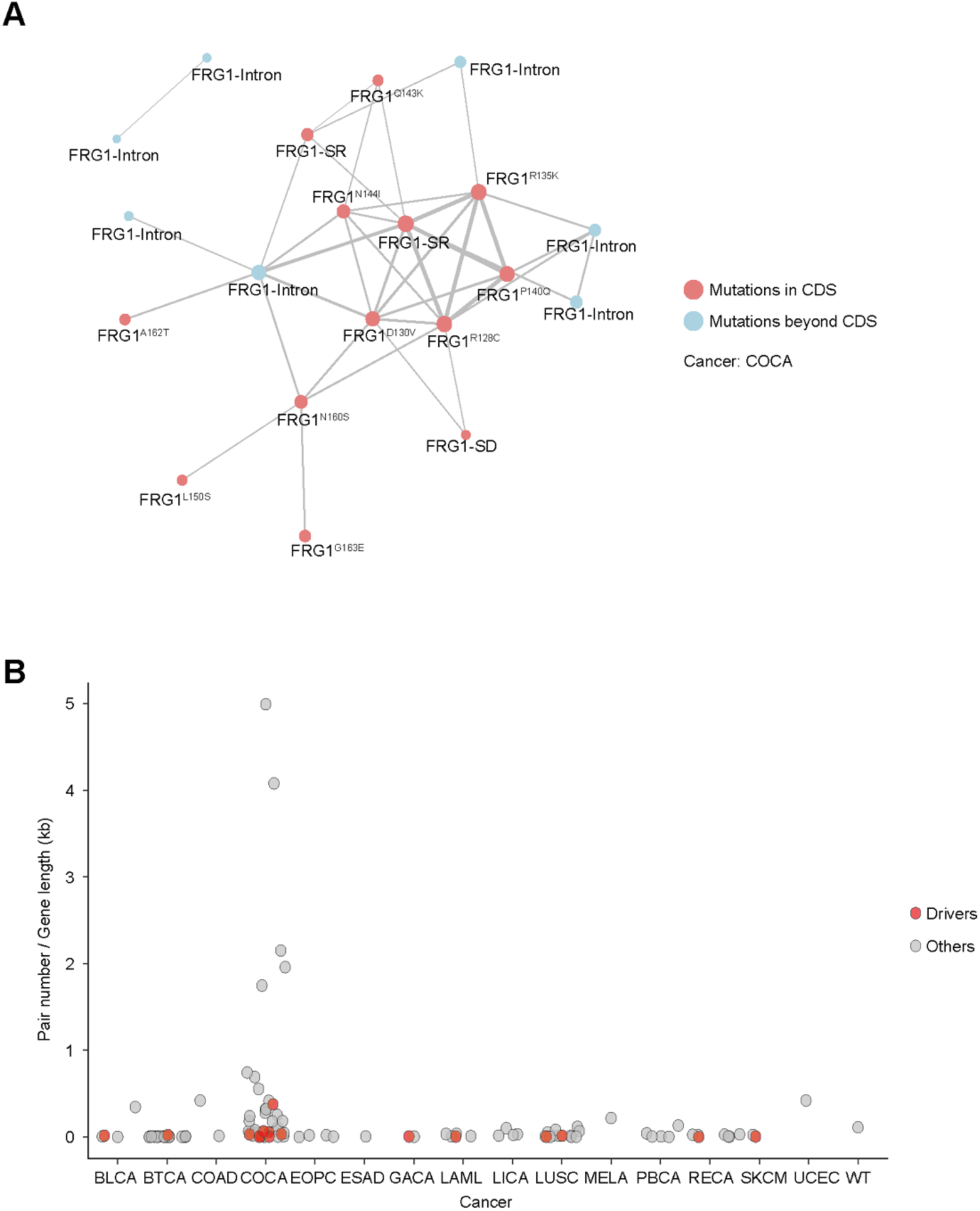
Intra-gene pairs. (A) Schematic showing the co-mutation pairs identified in *FRG1* in COCA, the colors stand for different mutation types, the dot size stands for the individual prevalence of a spefific mutation and the thickness of the line stands for the co-occurring prevalence. (B) Scatter plot showing the distribution of intra-gene pairs number normalized by gene length (kb) across cancer types. Each dot represents a given gene identified in the corresponding cancer types. The dots colored in red represent cancer drivers in CGC.

**Figure S4.**
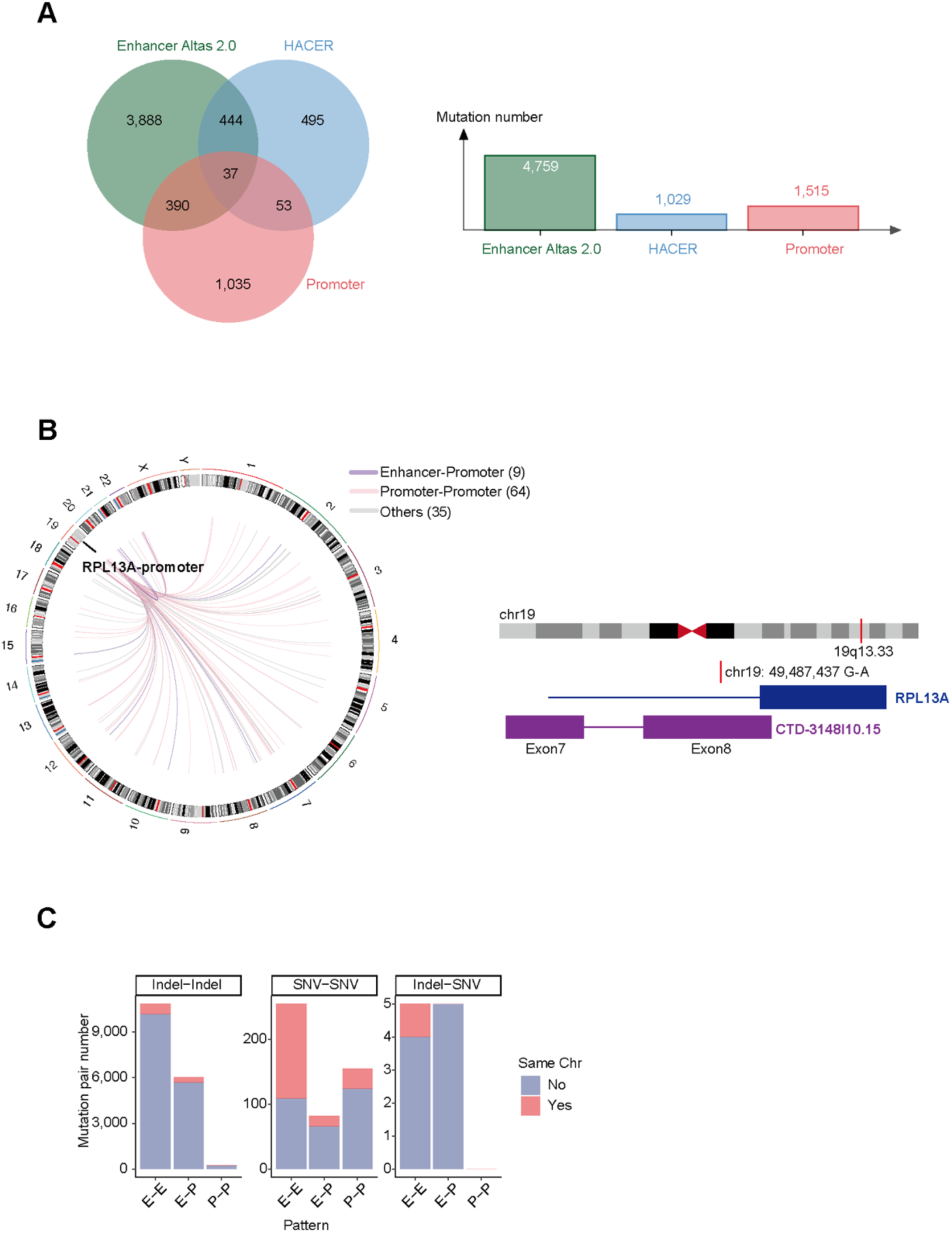
Non-coding co-mutation pairs. (A) Venn diagram of mutation number located in enhancer and promoter regions. (B) Chromatin looping diagram showing mutation pairs with *RPL13A* promoter mutation (MU38001837) in MELA. The colors of lines represent the EP patterns of mutation pairs. (C) The number of mutation pairs in each pattern classified by the mutation types and whether the two mutations are in the same chromosome.

**Figure S5.**
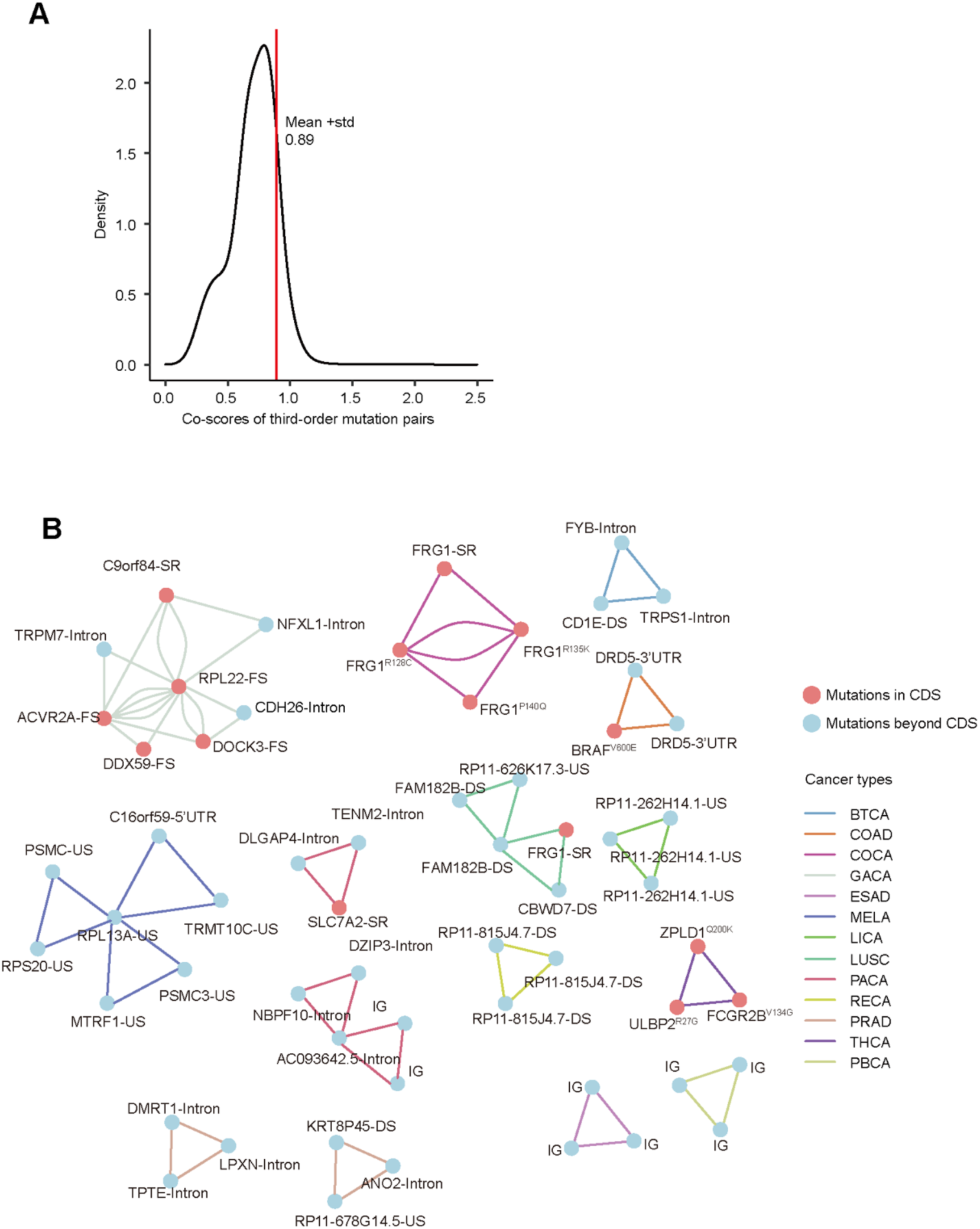
Third-order mutation pairs. (A) The co-score density of third-order mutation pairs. The mean plus standard deviation was about 0.89. To enhance the clarity of the image, the score range was set from 0 to 2.5, as there were only 155 mutation pairs with scores higher than 2.5. (B) Third-order mutation pairs with the highest frequency in different cancer types. DS: gene downstream variant, US: gene upstream variant, FS: frameshift variant, IG: intergenic variant.

## Method details

### ICGC data pre-processing

The original clinical dataset we used was simple somatic mutations (SSMs) recorded in ICGC database (https://docs.icgc-argo.org/docs/data-access/icgc-25k-data). We downloaded the mutation file for each cancer project in ICGC and merged the data with same cancer type but from different countries. We used LiftOver to convert the chromosomal positions of SSMs from the hg19 version to the hg38 version for subsequent data processing. Data filtering was performed in two steps: (1) Filtering mutations in repeat regions. The bed file for repeat regions was downloaded from the UCSC database, selecting the hg38 version of RepeatMasker under the Table Browser section (https://genome.ucsc.edu/cgi-bin/hgTables). The filtering was performed using the intersect function of bedtools^33^. (2) Excluding cancers with fewer than 100 donors. A sufficient sample size is essential for a reliable detection of co-occurring mutations. Cancer types with fewer than 100 donors were excluded to ensure statistical power. After these two filtering steps, a total of 45 cancer types, 19,152 patients, and 36,624,683 SSMs remained. Then we reorganized the mutation information in donor level for each cancer type and performed downstream calculation separately.

### FP-Growth algorithm

The FP-Growth (Frequent Pattern Growth) algorithm is a frequent pattern tree (FP-tree) structure-based data mining method used to extract frequent patterns from datasets without generating candidate item sets. The FP-Growth algorithm primarily includes the following steps^11^:

1. First scan to get supports for all items. These frequent items are ordered in descending frequency to create a list of frequent items. Items that do not reach the minimum support can be filtered in this step, because its superset will not meet the minimum support as well.
2. Second scan for FP-tree construction. Each transaction is processed in the order of frequent items and inserted into the FP-tree. If a subset of a transaction’s items already exists in the tree, their counters are incremented. New branches are added for new combinations of items. To illustrate, if *I* = {*i*_1_, *i*_2_, ⋯ , *i*_k_} represents a set of items in a transaction and *F* = {*f*_1_, *f*_2_, ⋯ , *f*_n_} is the set of frequent items, then the FP-tree is constructed using:

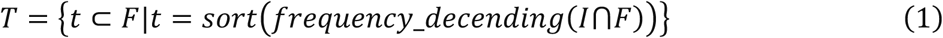

where each transaction *t* is added to the FP-tree based on the descending frequency of items in *F*.
3. Mining the FP-tree for frequency item sets. The search strategy is a partitioning-based, divide-and-conquer method. Item sets are extracted starting from the least frequent item, creating conditional bases for each item. A conditional FP-tree is then constructed for each item, and this process is recursively repeated to identify all frequent item sets.

In our research, mutation sets were defined as item sets for each donor, and support referred to the occurrence frequency of mutations. We used FP-Growth (https://borgelt.net/fpgrowth.html) to mine closed frequent item sets of mutations. The closed frequent itemset is a frequent itemset for which none of its immediate supersets have the same support as the itemset itself, providing the most complete information about patterns in the data without redundancy. Initially, we set the condition that mutations must co-occur in at least two donors, meaning the minimum support threshold was set as 2. We obtained second-order and third-order co-occurring mutation pairs by setting the itemset length as 2 and 3, respectively. Additionally, the FP-Growth algorithm was used to get frequency of each mutation by setting the itemset size as 1. Finally, the co-occurring frequency and individual frequencies of mutation pairs were organized using Python.

### Co-score calculation

We normalized the influence of donor and mutation counts across different cancers by calculating the co-score, which measures the co-occurrence degree of mutation pairs. The calculation method is described as Formula 2. Here, X represents the number of donors in which the mutation pair co-occur, N is the total number of donors in the cancer type, and M is the average number of mutations per donor in that cancer type. N and M are negatively correlated with the co-score. 1:1 coefficient was used for normalization, since the ranges of log_10_(N) and log_10_(M) were similar (Figures S1B and S1C). Based on the distribution of co-scores, we set the threshold as the mean plus standard deviation. The threshold for second-order mutation pairs was set to 0.97, and for third-order mutation pairs, it was set to 0.89. The mutation pairs with co-scores above the thresholds were retained as pairs with significantly co-occurring tendency.

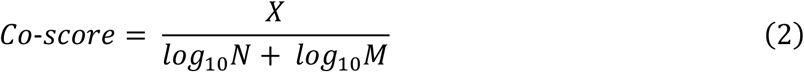

### Mutation pairs filtering

We classified mutations into two categories: single nucleotide variants (SNVs) and insertions/deletions (Indels). Based on this classification, we identified three types of mutation pairs: Indel-Indel, Indel-SNV, and SNV-SNV. To filter false positive mutation pairs in the second-order results, we excluded mutation pairs that met the following criteria: (1) SNV-SNV pairs that exhibit characteristics of the substrates of APOBEC3 cytosine deaminase enzymes, including: reference alleles being C or G with the “TCN” or “NGA” context, located on the same chromosome with a distance of less than or equal to 1,000 bp^15^; (2) Indel-SNV mutation pairs with a chromosomal distance of 3 bp or less, as these were likely the result of annotation errors^34^. After applying these filters, 4 SNV-SNV pairs and 3 Indel-SNV pairs were excluded, leaving a total of 101,007 second-order mutation pairs. For the third-order mutation pairs, we removed pairs where two mutations met the above criteria. This process resulted in the exclusion of 7 mutation pairs, leaving 18,366 third-order mutation pairs.

### Fisher’s exact test of mutation pairs

Fisher’s exact test is a statistical method used to determine if there are nonrandom associations between two categorical variables in a contingency table. We constructed the contingency table for each second-order mutation pairs by Python. As shown in the Table1, “M^+^” and “M^−^” represent the presence and absence of mutations, respectively, while *a*, *b*, *c* and *d* represent the number of cancer donors. For example, *a* represents the number of donors who carry both mutation 1 and mutation 2.

Based on Table 1, conditional on the margins of the table, *a* is distributed as a hypergeometric distribution with *a* + *c* draws from a population with *a* + *b* successes and c + d failures. The probability of obtaining such set of values is given by:

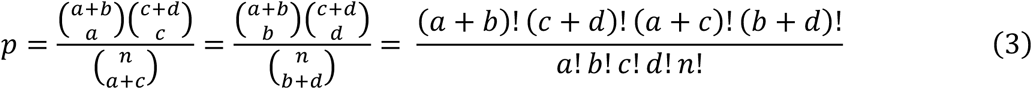

**Table1.** Mutation frequency contingency table.

|  | M<sub>1</sub><sup>+</sup> | M<sub>1</sub><sup>-</sup> |
| --- | --- | --- |
| M<sub>2</sub><sup>+</sup> | $a$ | $b$ |
| M<sub>2</sub><sup>-</sup> | $c$ | $d$ |

where 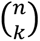 is the binomial coefficient, the symbol ! indicates the factorial operator, and *n* = *a* + *b* + *c* + *d.* We used the *scipy.stats* module in Python to perform one-sided Fisher’s exact test for each second-order mutation pairs. To reduce the occurrence of false positives, we applied Benjamini-Hochberg multiple test to correct the results of cancers with more than 100 mutation pairs, obtaining *q*-values. For the remaining cancers, we retained the original *p*-values. We then selected a significance threshold of 0.05. The mutation pairs with *p*-values or *q*-values less than the threshold were retained. Ultimately, we identified 100,933 significant second-order mutation pairs across 22 cancers.

### Jaccard index calculation

The Jaccard index (*J*) is used to gauge the similarity and diversity between sample sets. J is defined as the size of the intersection divided by the size of the union of the sample sets (Formula 4). In our research, we use *J* to define dependency of mutation pairs. After organizing the contingency table (Table 1), we use the Formula 5 to calculate *J*.

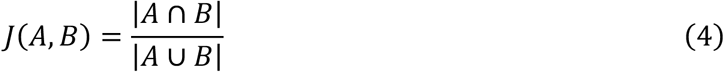

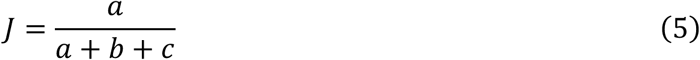

### Re-annotation of mutation consequence

ICGC provided multiple consequences for some mutations with the consideration of alternative splicing induced isoforms or two genes with very close genomic distance, leading to unwanted complexity in some cases. To facilitate our analysis in a global manner, we reannotated the consequence of each mutation in our results with the following two main steps. First, for the mutations located within gene body regions, we chose the consequence of canonical transcript according to MANE^35^ as its final annotation. Second, for the mutations beyond gene body or fail to match any of the canonical transcripts, we prioritized the functionality of consequences based on priori knowledge, which followed:

*[’missense_variant’,’synonymous_variant’,’frameshift_variant’,’splice_region_variant’,’splice_accept or_variant’,’splice_donor_variant’,’upstream_gene_variant’,’downstream_gene_variant’,’5_prime_U TR_variant’,’3_prime_UTR_variant’,’disruptive_inframe_deletion’,’disruptive_inframe_insertion’,’sto p_gained’,’5_prime_UTR_premature_start_codon_gain_variant’,’intron_variant’,’intergenic_region’,’ intragenic_variant’,’exon_variant’]*.

After the above steps, each mutation in our results got a unique annotation with was more probable to represent its functional type.

### ESM-IF1 scores of intra-gene mutation pairs mediated by missense mutations

ESM-IF1^25^ is a protein inverse folding model that can be used to assess the impact of sequence variations on protein structural stability. Missense mutations lead to changes in the protein sequence and may affect protein stability. Therefore, we can assess the synergistic effects of mutations on protein stability by comparing the stability of proteins with two mutations to that of proteins with a single mutation. We extracted mutation pairs within the same gene and both of which were missense mutations. The Uniprot ID and sequence for each protein were obtained from Uniprot (https://www.uniprot.org/id-mapping), and the protein structure was retrieved from AlphafoldDB^36,37^ based on the Uniprot ID. The wild-type, single-mutant, double-mutant protein sequences and wild-type structures were used as input, the script used to score the conditional log-likelihoods of the sequences for the corresponding structure was score_log_likelihoods.py (https://github.com/facebookresearch/esm/tree/main/examples/inverse_folding). Ultimately, we scored 44 missense mutation pairs from 22 proteins. The change in log-likelihood (delta log-likelihood) was calculated by subtracting the wild-type log-likelihood from the mutant log-likelihood. Additionally, the difference between the delta log-likelihood of the double-mutant and the sum of the delta log-likelihoods for the two single-mutants was calculated to evaluate the synergistic effect of combinational mutations on protein stability.

### De novo motif discovery of adjacent mutation pairs

We defined mutation pairs located within 20 nucleotides on the chromosome as adjacent mutation pairs, and those with greater distance as distant mutation pairs. We extracted the sequences between adjacent mutations, along with the flanking 10 base pairs on both sides by bedtools, and used MEME^38^ to search for potential motifs. Subsequently, for significant motifs, we used Tomtom^39^ to predict possible binding transcription factor proteins.

### Enhancer and promoter annotation

We used the enhancer databases EnhancerAtlas 2.0^40^ (http://enhanceratlas.org/) and HACER^41^ (https://bioinfo.vanderbilt.edu/AE/HACER/), which cover the most abundant cell lines from different sources. We selected the union of cell lines that correspond to the cancer tissues in our results, with detailed cell line information provided in Table S3. Data for the corresponding cell lines were downloaded from the two databases, respectively. The genomic data of enhancers was then converted from hg19 to hg38 using LiftOver, and mutations were annotated based on their genomic positions. A total of 4,759 mutations were located in enhancer regions annotated by EnhancerAtlas 2.0, and 1,029 mutations were located in enhancer regions annotated by the HACER database. The union of these two databases yielded 5,307 mutations located in enhancer regions. Promoter regions were defined as the 2,000 bp upstream of genes in the human hg38 genome. The BED file for promoters was downloaded from the UCSC Table Browser section (https://genome.ucsc.edu/cgi-bin/hgTables). In total, 1,515 mutations were annotated in promoter regions. There were 480 overlapping mutations between the enhancer and promoter regions, whose annotations were prioritized by E-P (enhancer mutation with promoter mutation), followed by P-P (promoter mutation with promoter mutation), E-E (enhancer mutation with promoter mutation).

### Correlation calculation of mRNA expression from tumor donors

We downloaded the integrated mRNA TPM (Transcripts Per Million) data for all TCGA patients from the R package TCGAplot. For mutation pairs annotated as E-P, P-P and E-E patterns, if both mutations had targeted gene annotations, we extracted the mRNA expression data of tumor samples in corresponding tissue and calculated the Pearson correlation coefficient between the two genes. The correspondence between cancer types in TCGA and ICGC in our results was provided in Table S4.

## Data and code availability

All the results of second-order and third-order mutation pairs can be queried and downloaded at https://co-mutation.streamlit.app/. The website was constracted by the *streamlit* module in Python. The bioinformatics pipeline is available at https://github.com/xiongyf520/co-mutation_code. Databases included ICGC dataset (https://docs.icgc-argo.org/docs/data-access/icgc-25k-data), Cancer Gene Census (https://cancer.sanger.ac.uk/census), HACER (https://bioinfo.vanderbilt.edu/AE/HACER/), EnhancerAtlas 2.0 (http://enhanceratlas.org/), Alphafold2 protein structure database (https://ftp.ebi.ac.uk/pub/databases/alphafold/latest/UP000005640_9606_HUMAN_v4.tar). Software and algorithms included FP-growth v6.20 (https://borgelt.net/fpgrowth.html), LiftOver (https://hgdownload.cse.ucsc.edu/admin/exe/linux.x86_64/liftOver), ESM-IF1 (https://github.com/facebookresearch/esm), bedtools v2.30.0, Python v3.9.12, R v4.2.1. Python modules included numpy 1.26.4, pandas 2.2.1, scipy 1.13.1. R packages included TCGAplot v8.0.0 (https://github.com/tjhwangxiong/TCGAplot), ggplot2 3.3.6, RCircos 1.2.2, ggbreak 0.1.2, dplyr 1.2.1.

## Acknowledgements

W.W. received funding from the National Science Foundation of China (82341207 and 31930016), the Peking-Tsinghua Center for Life Sciences, and Changping Laboratory. We express our gratitude to the Dr. X.M. at School of Electronics Engineering and Computer Science in Peking University for support in FP-Growth application, and Dr. Q.P., W.T., W.Z. and WJ.Z. in the lab for idea discussion. Our work is supported by High-performance Computing Platform of Peking University.

## Author contributions

The work presented in this paper was a collaborative effort of all authors. W.W. and Y.L. conceived and supervised the project. W.W., Y.L., Y.X. developed the bioinformatics pipeline and drafted the manuscript.

## Declaration of interests

W.W. is the founder and scientific advisor of EdiGene and Therorna. Authors declare that they have no other competing interests.

